# The grape powdery mildew resistance loci *Ren2, Ren3, Ren4D, Ren4U, Run1, Run1.2b, Run2.1*, and *Run2.2* activate different transcriptional responses to *Erysiphe necator*

**DOI:** 10.1101/2022.11.07.515491

**Authors:** Mélanie Massonnet, Summaira Riaz, Dániel Pap, Rosa Figueroa-Balderas, M. Andrew Walker, Dario Cantu

## Abstract

Multiple grape powdery mildew (PM) genetic resistance (*R*) loci have been found in wild grape species. Little is known about the defense responses associated with each *R* locus. In this study, we compare the defense mechanisms associated with PM resistance in interspecific crosses segregating for a single *R* locus from *Muscadinia rotundifolia* (*Run1, Run1.2b, Run2.1, Run2.2*), *Vitis cinerea* (*Ren2*), *V. romanetii* (*Ren4D* and *Ren4U*), and the interspecific hybrid Villard blanc (*Ren3*). By combining optical microscopy, visual scoring, and biomass estimation, we show that the eight *R* loci confer resistance by limiting infection at different stages. We assessed the defense mechanisms triggered in response to PM at 1 and 5 days post inoculation (dpi) via RNA sequencing. To account for the genetic differences between species, we developed for each accession a diploid synthetic reference transcriptome by incorporating into the PN40024 reference homozygous and heterozygous sequence variants and *de novo* assembled transcripts. Most of the *R* loci exhibited a higher number of differentially expressed genes (DEGs) associated with PM resistance at 1 dpi compared to 5 dpi, suggesting that PM resistance is mostly associated with an early transcriptional reprogramming. Comparison of the PM resistance-associated DEGs showed a limited overlap between pairs of *R* loci, and nearly half of the DEGs were specific to a single *R* locus. The largest overlap of PM resistance-associated DEGs was found between *Ren3*^+^, *Ren4D*^+^, and *Ren4U*^+^ genotypes at 1 dpi, and between *Ren4U*^+^ and *Run1*^+^ accessions at 5 dpi. The *Ren3*^+^, *Ren4D*^+^, and *Ren4U*^+^ were also found to have the highest number of *R* locus-specific DEGs in response to PM. Both shared and *R* locus-specific DEGs included genes from different defense-related categories, indicating that the presence of *E. necator* triggered distinct transcriptional responses in the eight *R* loci.

## Introduction

Most cultivated grapevines (*Vitis vinifera* ssp. *vinifera*) are highly susceptible to powdery mildew (PM), a disease caused by the ascomycete *Erysiphe necator* Schwein (syn. *Uncinula necator*). *Erysiphe necator* is an obligate biotrophic pathogen that can infect any green tissue of the host. PM infections lead to reduced yield, and impaired fruit composition and wine quality (Calonnec *et al*., 2004; Stummer *et al*., 2005). Genetic resistance to PM has been found in several wild grapes, such as the North American grapes *Muscadinia rotundifolia* and *Vitis cinerea*, the Chinese *Vitis* species *V. piasezkii, V. romanetii, V. quinquangularis* and *V. pseudoreticulata*, and some Central Asian accessions of *V. vinifera* ssp. *sylvestris* (Dry *et al*., 2019). So far, thirteen PM resistance (*R*) loci have been genetically mapped (Dry *et al*., 2019; Karn *et al*., 2021) and named *Resistance to Uncinula necator* (*Run*) or *Resistance to Erysiphe necator* (*Ren*). Different allelic forms have been found for some of *Run* and *Ren* loci (Dry *et al*., 2019; Massonnet *et al*., 2022).

For most PM-resistant grape accessions, resistance to *E. necator* is associated with a programmed cell death (PCD)-mediated response (Qiu *et al*., 2015; Dry *et al*., 2019). Because PCD occurs in epidermal cells post-penetration of *E. necator*, the recognition of *E. necator*’s effectors by intracellular nucleotide-binding leucine-rich repeat (NLR) proteins is likely the trigger of PCD (Qiu *et al*., 2015; Dry *et al*., 2019). Only one NLR gene associated with PM resistance, *MrRUN1*, has been characterized in *M. rotundifolia* G52 (Feechan *et al*., 2013) and candidate NLR genes associated with *Run1.2b* and *Run2.2* have been proposed for *M. rotundifolia* Trayshed (Massonnet *et al*., 2022).

In plants, NLR activation leads to multiple cellular responses, such as the generation of reactive oxygen species (ROS), calcium oscillations, kinase activation, and an extensive transcriptional reprogramming resulting in the activation of defense-related mechanisms, including the biosynthesis of pathogenesis-related (PR) proteins and antimicrobial compounds, and cell wall modifications (Dangl *et al*., 2013; Lolle *et al*., 2020). NLR-triggered immunity generally involves PCD at the site of infection, which inhibits the development of the pathogen (Dangl *et al*., 2013; Lolle *et al*., 2020). In grapes, few studies have investigated the transcriptomic responses to *E. necator* infection in PM-resistant accessions (Amrine *et al*., 2015; Jiao *et al*., 2021; Weng *et al*., 2014). For example, comparative transcriptomics revealed that *E. necator* infection leads to diverse whole-genome transcriptional responses among Central Asian *V. vinifera* accessions carrying different allelic forms of the *Ren1* locus (Amrine *et al*., 2015). Whether the resistance conferred by different *R* loci depends on the activation of the same or different defense responses is still an open question. Understanding the functional differences and overlap between *R* loci will help select the most functionally diverse *R* loci to develop new *V. vinifera* cultivars that combine durable resistance to PM and high-quality fruit production (Michelmore *et al*., 2013). However, comparing genome-wide transcriptional changes in response to PM between *R* loci from different *Vitis* species is challenging. Comparative transcriptomics using RNA sequencing (RNA-seq) data relies on a reference transcriptome to evaluate transcript abundance. Recent studies showed that grape genomes are highly heterozygous and substantially differ between *Vitis* species (Zhou *et al*., 2019; Liang *et al*., 2019; Cochetel *et al*., 2021; Minio *et al*., 2022). A single haploid reference transcriptome from a PM-susceptible *V. vinifera* biases the alignment of the RNA-seq reads, underestimating the expression of alleles that are less similar to the reference, and confounding the subsequent testing of differential gene expression.

In this study, we aimed to identify the defense mechanisms associated with eight *R* loci (*Ren2, Ren3, Ren4D, Ren4U, Run1, Run1.2b, Run2.1*, and *Run2.2*) and to evaluate the functional differences and overlap between them. First, we monitored PM disease development on leaves of eight PM-resistant breeding lines, each representing a genetic *R* locus, as well as four PM-susceptible sib lines and two PM-susceptible *V. vinifera* parents. Defense mechanisms associated with each *R* locus were then assessed by profiling the leaf transcriptome with or without *E. necator* inoculations. To enable the comparison of the transcriptional modulation in response to PM, we constructed comparable reference transcriptomes incorporating both sequence variant information into the reference transcriptome of PN40024 and *de novo* assembled transcripts. We also refined the functional annotation of PN40024 predicted proteome to focus our analysis on genes involved in defense mechanisms. Defense-related genes differentially expressed in response to the pathogen were first compared between PM-resistant and PM-susceptible accessions to identify the genes with a transcriptional modulation associated with PM resistance. The latter ones were then compared between PM-resistant genotypes to evaluate the functional overlap between the different *R* loci.

## Materials and Methods

### Plant material and evaluation of PM development

Fourteen grape accessions were used in this study: eight carrying one *R* locus, six sib lines without any *R* locus, and two susceptible *V. vinifera* parents. Information about the pedigree of each accession is provided in **Supplementary Table 1**.

PM susceptibility was evaluated with a detached leaf assay as described in Pap *et al*. (2016). Detached leaves were stained with Coomassie Brilliant Blue R-250 at 5 days post inoculation (dpi) as in Riaz *et al*. (2013). Visual disease susceptibility scores at 14 dpi were compared using a Kruskal-Wallis test followed by a post hoc Dunn’s test (*P* ≤ 0.05).

For transcriptional profiling, we followed the inoculation protocol described in Amrine *et al*. (2015). For each accession three plants were inoculated with *E. necator* C-strain (Jones et al., 2014) and three plants were mock-inoculated. Two leaves from each plant were collected 1 and 5 dpi and immediately frozen in liquid nitrogen. Leaves from an individual plant were pooled together and constitute a biological replicate. Three biological replications were obtained for each treatment.

### RNA extraction, library preparation, and sequencing

RNA extraction and library preparation were performed as in Amrine *et al*. (2015). cDNA libraries were sequenced using Illumina HiSeq2500 and HiSeq4000 sequencers (DNA Technologies Core, University of California, Davis, CA, USA) as 50-bp single-end reads (**Supplementary Table 2**). Sequencing reads of the accessions e6-23 (*Run1.2b*^+^) and 08391-29 (*Run2.2*^+^) were retrieved from the NCBI BioProject PRJNA780568.

### Reconstruction of accession-specific reference transcriptomes

For each accession, we constructed a reference transcriptome composed of a diploid synthetic transcriptome and *de novo* assembled transcripts. The diploid synthetic transcriptome was reconstructed using sequence variant information. First, adapter sequences were removed and RNA-seq reads were filtered based on their quality using Trimmomatic v.0.36 (Bolger *et al*., 2014) and these settings: LEADING:3 TRAILING:3 SLIDINGWINDOW:10:20 MINLEN:20. Quality-trimmed reads of the 12 samples from each accession were concatenated into a single file and mapped onto a combined reference genome composed of *V. vinifera* PN40024 V1 (Jaillon *et al*., 2007) and *E. necator* C-strain genome (Jones *et al*., 2014) following the STAR 2-pass mapping protocol (v.2.5.3a; Dobin *et al*., 2013; Engström *et al*., 2013). PCR and optical duplicates were removed with Picard tools (v.2.0.1 http://broadinstitute.github.io/picard/), and reads were split into exon segments using SplitNCigarReads from GATK v.3.5-0-g36282e4 (McKenna *et al*., 2010). GATK HaplotypeCaller was used to call sequence variants with the following parameters: -ploidy 2 -stand_call_conf 20.0 -stand_emit_conf 20.0 -dontUseSoftClippedBases. Variants were filtered using GATK VariantFiltration with these settings: -window 35 -cluster 3 -filterName FS -filter “FS > 30.0” -filterName QD -filter “QD < 2.0”. Variants passing all filters were selected using GATK SelectVariants with “--excludeFiltered” parameter. On average, 265,900 ± 39,570 variants were detected per genotype (**Supplementary Table 3**). Variants in grape protein-coding regions were extracted using bedtools intersect v.2.19.1 (Quinlan, 2014). Two separate genomes were reconstructed for each genotype using the vcf-consensus tool from vcftools (Danecek *et al*., 2011). The first genome was reconstructed by incorporating both homozygous and heterozygous alternative 1 (ALT1) variants relative to the PN40024 genome, while the second genome was reconstructed using both homozygous and heterozygous alternative 2 (ALT2) variants. A new annotation file was created for each genome from its corresponding variant information. CDS were extracted using gffread from Cufflinks v.2.2.1 (Trapnell *et al*., 2010).

For each genotype, quality-trimmed reads were mapped onto their respective diploid synthetic grape transcriptome and *E. necator* C-strain CDS (Jones *et al*., 2014) using Bowtie2 v.2.3.4.1 (Langmead and Salzberg, 2012) and the parameters --end-to-end --sensitive --un. For each grape accession, *de novo* assembly was performed using unmapped reads from the 12 RNA-seq libraries and TRINITY v.2.4.0 (Grabherr *et al*., 2011). A total of 509,960 sequences were reconstructed, corresponding to 36,426 ± 2,585 sequences per accession (**Supplementary Table 4**). To reduce sequence redundancy, reconstructed transcripts from all 14 genotypes were clustered using CD-HIT-EST v.4.6.8 (Li and Godzik, 2006) and an identity threshold of 90%. The longest representative sequence of each of the 98,340 transcript clusters was used as input for TransDecoder v.3.0.1 (https://github.com/TransDecoder/TransDecoder). For the 2,174 transcripts with an open reading frame protein containing a start and a stop codon, the longest CDS was selected. CDS redundancy was reduced by clustering using CD-HIT-EST v.4.6.8 (Li and Godzik, 2006), with coverage and identity thresholds of 100%. In total, 2,070 CDS were retained (**Supplementary Data 1**). Taxonomic analysis of the predicted proteins was performed using Megan v.6.12.5 (Huson *et al*., 2016) with default parameters after aligning predicted peptides against the RefSeq protein database (ftp://ftp.ncbi.nlm.nih.gov/refseq, retrieved 17 January 2017). The 103 peptides assigned to proteobacteria, opisthokonts, and viruses were considered microbial contaminants; the remaining 1,967 peptides were assigned to grape (**Supplementary Table 5**).

To evaluate the effect of the reference transcriptome on the mapping rate (*i.e*. the percentage of RNA-seq reads aligning onto the reference transcriptome), quality-trimmed RNA-seq reads were aligned onto two reference transcriptomes: (i) the combined protein-coding sequences of *V. vinifera* PN40024 V1 (Jaillon *et al*., 2007) and *E. necator* C-strain (Jones *et al*., 2014), (ii) the combined accession-specific diploid synthetic grape transcriptome, the *de novo* assembled CDS, and *E. necator* C-strain CDS (Jones *et al*., 2014), using Bowtie2 v.2.3.4.1 (Langmead and Salzberg, 2012) and the parameters --end-to-end --sensitive --un. For each accession, the difference of mapping rates between the two reference transcriptomes were tested using Kruskal-Wallis test followed by post hoc Dunn’s test (*P* ≤ 0.05).

### Gene expression analysis

Transcript abundance was assessed using Salmon v.1.5.1 (Patro *et al*., 2017) and these parameters: --gcBias --seqBias --validateMappings. For each accession, a transcriptome index file was built using the accession’s diploid synthetic transcriptome combined with the *de novo* assembled transcripts and the *E. necator* C-strain transcriptome, PN40024 V1 and *E. necator* C-strain genomes as decoys, and a k-mer size of 13. Read counts were computed using the R package tximport v.1.20.0 (Soneson *et al*., 2015). Read counts of the two haplotypes of each gene locus of the diploid synthetic transcriptome were combined. Read-count normalization of the grape transcripts and statistical testing of differential expression were performed using DESeq2 v.1.16.1 (Love *et al*., 2014). The sample 14305-001_H_5dpi_1 was removed from the RNA-seq analysis because of its low mapping rate (**Supplementary Table 2**).

### Functional annotation of the defense-related genes

To determine the defense mechanisms triggered by the presence of *E. necator*, we refined the functional annotation of the grape predicted proteins involved in the following processes: pathogen recognition by receptor-like kinases (RLKs) and intracellular receptors (NLRs), ROS production and scavenging, nitric oxide (NO) production, calcium oscillations, MAPK cascade, salicylic acid (SA), jasmonic acid (JA), ethylene (ET), and abscisic acid (ABA) signaling, pathogenesis-related (PR) protein and phytoalexin biosynthesis, and cell wall reinforcement. Functional annotation of the grape predicted proteins was assigned based on sequence homology with *Arabidopsis thaliana* predicted proteins involved in the aforementioned functional categories and protein domain composition. Grape proteins were aligned onto *A. thaliana* proteins (Araport11_genes.201606.pep.fasta; https://www.arabidopsis.org/download/index.jsp) using BLASTP v.2.6.0. Alignments with an identity greater than 30% and a reciprocal reference:query coverage between 75% and 125% were kept. For each grape protein, the alignment with the highest product of identity, query coverage, and reference coverage was selected to determine a homologous protein in *A. thaliana*. We verified that each grape protein and its assigned *A. thaliana* had a similar domain composition. Grape and *Arabidopsis thaliana* predicted proteins were scanned with hmmsearch from HMMER v.3.3.1 (http://hmmer.org/) and the Pfam-A Hidden Markov Models (HMM) database (El-Gebali *et al*., 2019; downloaded on 29 January 2021). Protein domains with an independent E-value less than 1.0 and an alignment covering at least 50% of the HMM were selected. Grape predicted proteins having a similar domain composition than their *A. thaliana* homologues were retained. The functional annotation of defense-related genes used in this study can be found in **Supplementary Table 6**.

## Results

### Powdery mildew resistance loci exhibit different intensity and timing of response to *E. necator*

PM disease severity was evaluated on detached leaves from fourteen grape accessions, including twelve interspecific accessions, and two PM-susceptible *V. vinifera*, Malaga Rosada and F2-35 (**Table 1, Supplementary Table 1**). The twelve interspecific hybrids included four pairs of siblings derived from backcrosses with one of the following resistant accessions: *V. romanetii* C166-043 (*Ren4D*^+^), *V. romanetii* C166-026 (*Ren4U*^+^), *M. rotundifolia* G52 (*Run1*^+^), *M. rotundifolia* Trayshed (*Run2.2*^+^) (Ramming *et al*., 2011; Riaz *et al*., 2011; Feechan *et al*., 2013). The four remaining interspecific accessions were 09390-023 (*Ren2*^+^), 07712-06 (*Ren3*^+^), e6-23 (*Run1.2b*^+^), and 09705-45 (*Run2.1*^+^); these inherited PM resistance from *V. cinerea* B9, the interspecific hybrid Villard blanc, *M. rotundifolia* Trayshed, and *M. rotundifolia* Magnolia, respectively (Dalbó e*t al*., 2001; Riaz e*t al*., 2011; Zyprian e*t al*., 2016).

**Table 1:** Description of the fourteen grape genotypes used in this study. Details about their pedigree are provided in **Supplementary Table 1**. Siblings deriving from the same cross are indicated with an asterisk. BC, backcross.

| Accession name | *R* locus | *R* locus origin | BC level | % *V. vinifera* |
| --- | --- | --- | --- | --- |
| 09390-023 | *Ren2*<sup>+</sup> | *V. cinerea* B9 | F1 | 50 |
| 07712-06 | *Ren3*<sup>+</sup> | Villard blanc | Complex | 78 |
| 13353-55* | *Ren4D*<sup>+</sup> | *V. romanetii* C166-043 | BC2 | 87.5 |
| 13353-33* | *Ren4D*<sup>-</sup> | - | BC2 | 87.5 |
| 14305-002* | *Ren4U*<sup>+</sup> | *V. romanetii* C166-026 | Complex | 89.1 |
| 14305-001* | *Ren4U*<sup>-</sup> | - | Complex | 89.1 |
| 14375-059* | *Run1*<sup>+</sup> | *M. rotundifolia* G52 | BC4 | 96.9 |
| 14375-063* | *Run1*<sup>-</sup> | - | BC4 | 96.9 |
| e6-23 | *Run1.2b*<sup>+</sup> | *M. rotundifolia* Trayshed | BC2 | 87.5 |
| 09705-45 | *Run2.1*<sup>+</sup> | *M. rotundifolia* Magnolia | BC2 | 87.5 |
| 08391-029* | *Run2.2*<sup>+</sup> | *M. rotundifolia* Trayshed | BC3 | 93.8 |
| 08391-028* | *Run2.2*<sup>-</sup> | - | BC3 | 93.8 |
| F2-35 | - | - | - | 100 |
| Malaga Rosada | - | - | - | 100 |

At 5 dpi, extensive hyphal growth and conidiophores were observed on leaves from the two PM-susceptible *V. vinifera* cultivars, F2-35 and Malaga Rosada, and all the sib lines devoid of PM resistance loci (**Figure 1A**). Little or no hyphal growth was visible on the leaves of the accessions carrying *Ren4D, Ren4U, Run1*, and *Run1.2b* loci. Some secondary and tertiary hyphae were found on *Ren2*^+^, *Ren3*^+^, *Run2.1*^+^, and *Run2.2*^+^ genotypes (**Figure 1A**). PM infection was also assessed at an advanced disease development stage (14 dpi) using visual scoring (**Supplementary Figure 1A**). Infection rate at 5 and 14 dpi was similar for all genotypes except *Ren2*^+^ and *Ren3*^+^, which both had an extensive mycelium growth on their leaves at 14 dpi (Kruskal-Wallis test followed by post-hoc Dunn’s test; *P* ≤ 0.05).

**Figure 1:**
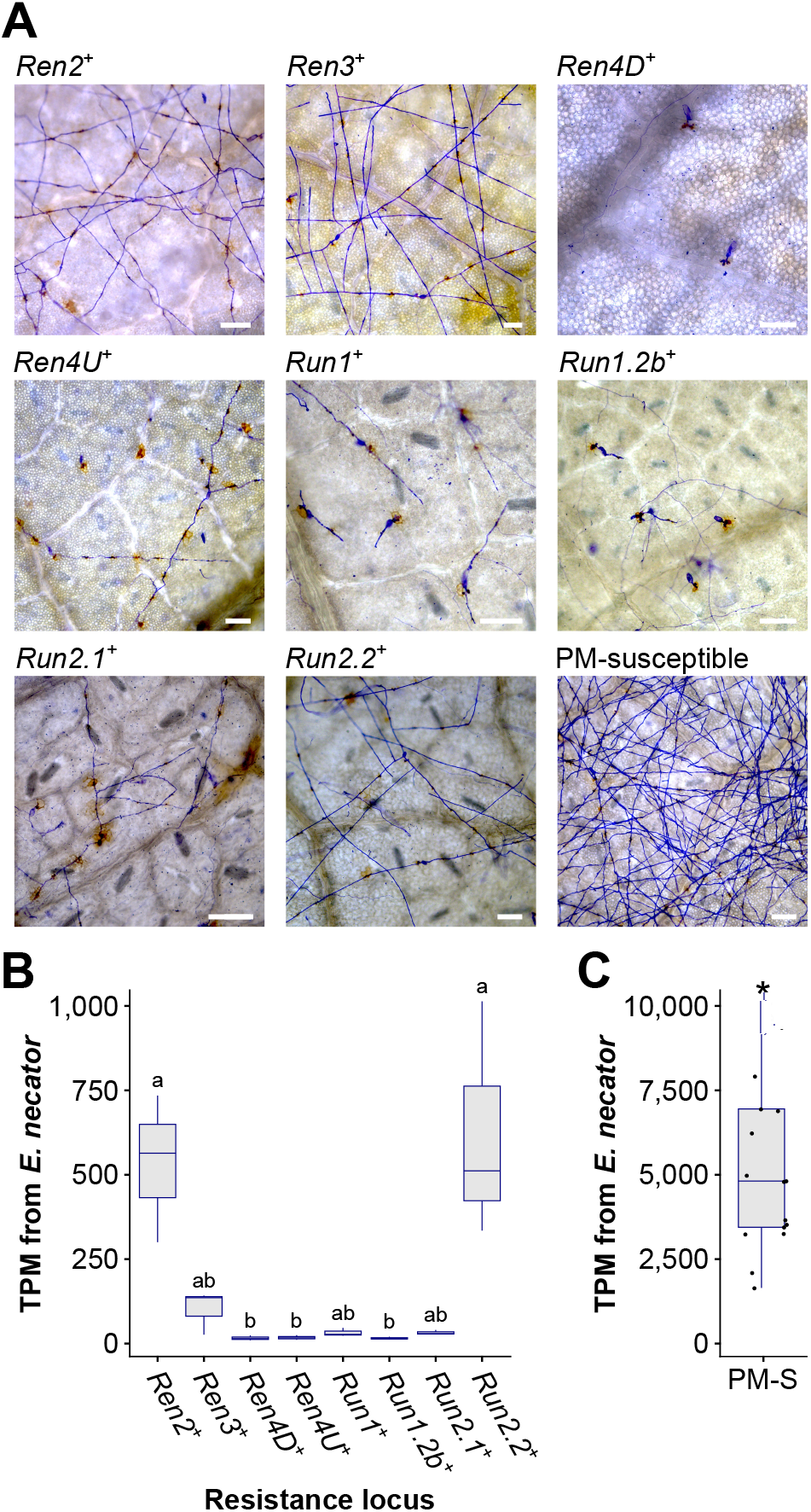
Powdery mildew disease development at 5 days post-inoculation. (**A**) Micrographs of detached leaves inoculated with *E. necator*. Scale = 100 μm. Total Transcripts per Million (TPM) derived from *E. necator* transcriptome in PM-resistant (**B**) and PM-susceptible (PM-S) accessions (**C**). Significant differences between PM resistance loci are indicated by different letters (Kruskal-Wallis test followed by post hoc Dunn’s test; *P* ≤ 0.05). Significant difference between grape accessions with and without a PM resistance locus is indicated by an asterisk (Kruskal-Wallis test, *P* = 4.0 × 10^−8^).

PM biomass on leaves was estimated by measuring *E. necator* transcript abundance at 5 dpi. As expected, significantly lower *E. necator* transcript counts were found in genotypes carrying a PM *R* locus (**Figure 1B,C**). Significant differences in *E. necator* transcripts were also observed between *Ren2*^+^, *Run2.2*^+^, and the other PM-resistant accessions (Kruskal-Wallis test followed by post hoc Dunn’s test; *P* ≤ 0.05). *E. necator* transcript abundance at 5 dpi and PM infection scores at 14 dpi correlated well (*R*^2^ = 0.71; **Supplementary Figure 1B**), which suggests that pathogen transcript abundance is a reliable measure of PM susceptibility.

These results show that all *R* loci confer resistance to PM in *V. vinifera*. However, PM resistance level varies between *R* loci, suggesting differences in the perception of the pathogen and/or in the efficiency of the defense responses.

### Construction of comparable reference transcriptomes

To determine if the differences in PM development between *R* loci were associated with differences in the defense mechanisms induced, we profiled the leaf transcriptomes of the fourteen genotypes at 1 and 5 dpi using RNA-seq. Mock-inoculated samples were collected as controls. Because the genotypes have diverse and distant genetic backgrounds, we built comparable reference transcriptomes for each genotype by incorporating sequence variant information into the predicted transcriptome of PN40024 and by adding *de novo* assembled transcripts that are not found among the annotated CDS of PN40024.

RNA-seq reads were first used to identify sequence polymorphisms between all sequenced transcriptomes and the PN40024 reference (Jaillon *et al*., 2007). On average, 109,645 ± 17,463 single-nucleotide polymorphisms (SNPs) and 1,208 ± 119 short insertion-deletions (INDELs) were detected in the protein-coding regions of each grape accession (**Figure 2**; **Supplementary Table 3**). Although the number of homozygous SNPs was quite stable across genotypes (35,490 ± 4,667), the number of heterozygous SNPs varied extensively, ranging from 54,154 in *V. vinifera* cv. Malaga Rosada to 116,032 in the *Ren2*^+^ accession (Malaga Rosada x *V. cinerea* B9; **Figure 2**). Comparison of the SNPs detected in the fourteen leaf transcriptomes compared to *V. vinifera* PN40024 protein-coding sequences distinguished the *Ren2*^+^ accession from the other grape accessions, likely because it is the only F1 individual in this study (**Table 1**; **Figure 2**). Siblings and respective *V. vinifera* parents clustered together confirming the validity of the variant calling approach (**Figure 2**). Analysis of genetic relatedness also confirmed sibling and parent-offspring relationships (**Supplementary Figure 2**). Homozygous and heterozygous variants of each grape genotype were incorporated into the CDS of PN40024 to produce a diploid reference transcriptome for each genotype (**Figure 3A**).

**Figure 2:**
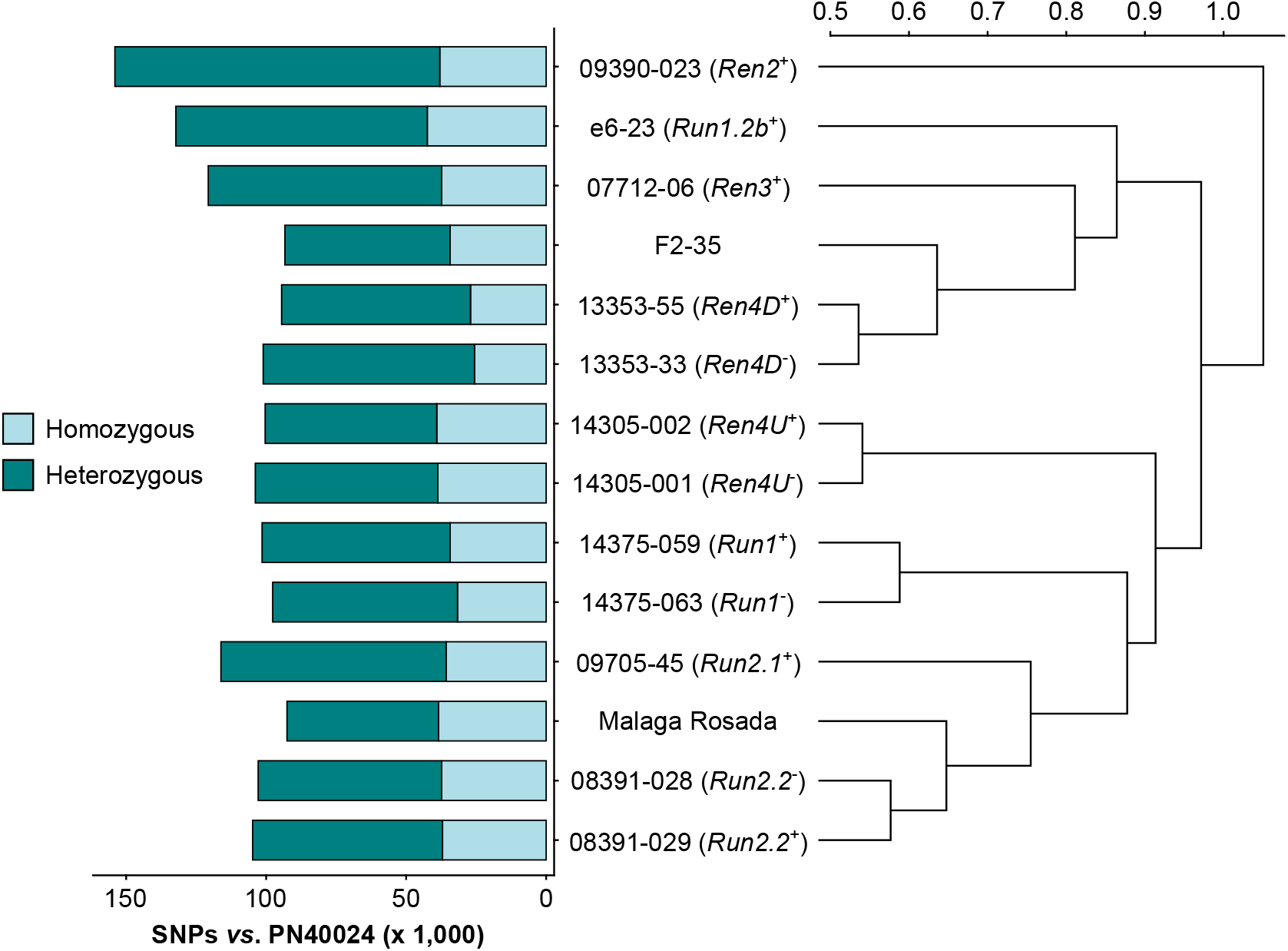
SNPs detected in the leaf transcriptome of the 14 grape accessions compared to the protein-coding sequences of *Vitis vinifera* PN40024. The Pearson correlation coefficients of the pairwise SNP comparison were converted into distance coefficients to define the height of the dendrogram branches.

**Figure 3:**
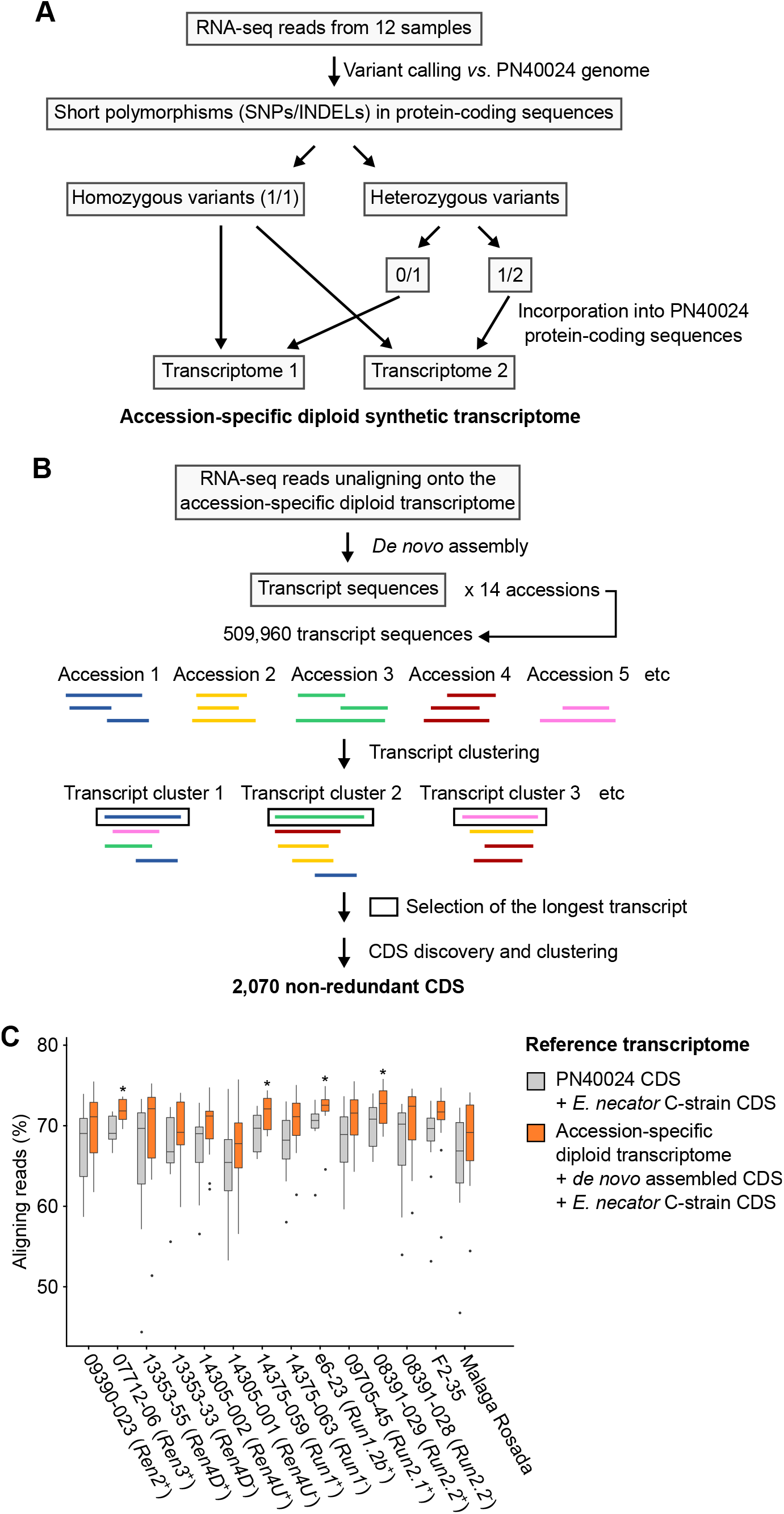
Adapting the grape transcriptome to each accession enhances the alignment of RNA-seq reads. Schematic representations of the bioinformatics approaches used to reconstruct the diploid synthetic transcriptome of each grapevine accession (**A**) and to cluster the *de novo* assembled transcripts (**B**). (**C**) Effect of the reference transcriptome on the percentage of aligning reads. For each genotype, difference of mapping rate with the combined transcriptomes of PN40024 and *E. necator* was tested using a Kruskal-Wallis test. * indicates *P* value ≤ 0.05.

The RNA-seq reads that did not align onto their respective diploid transcriptome reference were *de novo* assembled in order to construct transcripts that are not present in the PN40024 transcriptome (**Figure 3B**). All *de novo* assembled transcripts from the fourteen accessions were clustered to obtain a non-redundant dataset. After sequence clustering, CDS discovery, and removal of non-plant sequences, 1,967 non-redundant plant CDS were identified (See methods; **Supplementary Table 5**). Based on transcript clustering, each grape accession possessed on average 1,201 ± 31 of the grape *de novo* assembled CDS. As expected, siblings grouped together based on shared CDS (**Supplementary Figure 3**). Incorporating the sequence polymorphisms and *de novo* assembled CDS into the reference transcriptome increased the percentage of aligning RNA-seq reads by 2.5 ± 0.9% per sample (**Figure 3C**) and likely also increased the mapping specificity.

### Annotation of the grape defense-related genes

To determine the defense mechanisms involved in the response to *E. necator*, we refined the functional annotation of the grape predicted proteins based on protein domain composition and homology with *Arabidopsis thaliana* predicted proteins. We identified 2,694 grape genes involved in the following processes: pathogen recognition by receptor-like kinases (RLKs) and intracellular receptors (NLRs), ROS production and scavenging, nitric oxide (NO) production, calcium oscillations, MAPK cascade, salicylic acid (SA), jasmonic acid (JA), ethylene (ET), and abscisic acid (ABA) signaling pathways, pathogenesis-related (PR) protein and phytoalexin biosynthesis, and cell wall reinforcement (**Table 2**; **Supplementary Table 6**). These included 22 *de novo* assembled CDS, including nine NLRs, a glutaredoxin, three pathogenesis-related (PR) protein 14-like proteins, and three ethylene-responsive transcription factors (ERFs) (**Supplementary Table 6**).

**Table 2:** Number of defense-related genes identified among the predicted proteins of PN40024 CDS and the *de novo* assembled CDS. NLR, nucleotide-binding leucine-rich repeat protein.

| Defense-related functional category | PN40024 CDS | *de novo* CDS | Total |
| --- | --- | --- | --- |
| Receptor-like kinases (RLKs) | 450 | 1 | 451 |
| Nucleotide-binding leucine-rich repeat proteins (NLRs) | 309 | 9 | 318 |
| Calcium signaling | 109 | 0 | 109 |
| MAPK signaling | 83 | 0 | 83 |
| Reactive oxygen species (ROS) production | 106 | 0 | 106 |
| Reactive oxygen species (ROS) scavenging | 216 | 1 | 217 |
| Nitric oxide (NO) production | 8 | 0 | 8 |
| Salicylic acid-mediated signaling pathway | 26 | 0 | 26 |
| Jasmonic acid-mediated signaling pathway | 113 | 1 | 114 |
| Ethylene-mediated signaling pathway | 168 | 3 | 171 |
| Absciscic acid-mediated signaling pathway | 111 | 0 | 111 |
| Pathogenesis-related (PR) proteins | 374 | 3 | 377 |
| Phytoalexin biosynthesis | 339 | 1 | 340 |
| Cell wall reinforcement | 260 | 3 | 263 |
| **Total** | **2,672** | **22** | **2,694** |

### Assessment of the transcriptional modulations associated with powdery mildew resistance

PM- and mock-inoculated leaf transcriptomes were compared at 1 and 5 dpi to identify the defense-related genes that were differentially expressed in response to *E. necator* (**Figure 4**; **Supplementary Table 7**). In the PM-resistant vines *Ren3*^+^, *Ren4D*^+^, *Ren4U*^+^, and *Run1.2b*^+^, up- and down-regulated genes were more numerous at 1 dpi relative to 5 dpi, while only few differentially expressed genes (DEGs) were detected in *Run2.1*^+^ leaves. The opposite pattern was observed in the *Run2.2*^+^ accession as well as four of the five PM-susceptible plants. These results suggest that disease resistance is mostly associated with an early transcriptional reprogramming.

**Figure 4:**
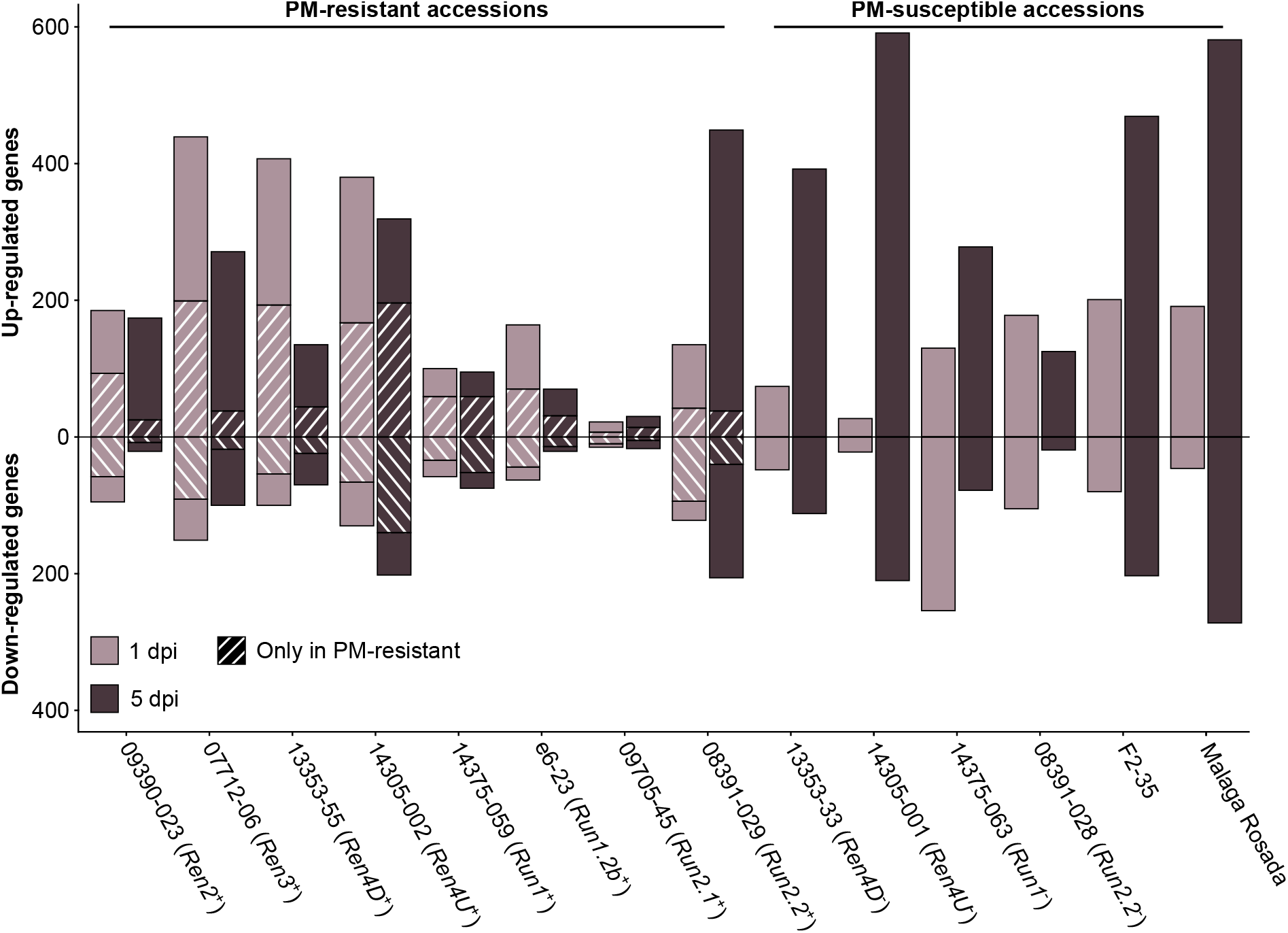
Defense-related genes differentially expressed in response to *E. necator* at 1 and 5 dpi. The genes uniquely differentially expressed by *E. necator* in the grape accessions possessing a *R* locus were identified by comparing the up- and down-regulated genes in PM-resistant leaves with the ones detected in PM-susceptible vines at each time point.

To determine the transcriptional modulations associated with PM resistance, defense-related DEGs in PM-resistant accessions were compared with the ones found in PM-susceptible individuals at each time point. On average, 39.7 % ± 16.6 % of the up-regulated genes and 52.7 % ± 19.0 % of the down-regulated genes detected in PM-resistant accessions were not found in PM-susceptible plants (**Figure 4**). PM resistance-associated DEGs were more numerous at 1 dpi compared to 5 dpi in all PM-resistant genotypes except the *Ren4U*^+^ vine. This supports the hypothesis that an early transcriptional reprogramming caused by the prompt perception of *E. necator* by intracellular NLR is crucial for PM resistance. However, the PM-resistant accessions exhibited different amounts of defense-related DEGs in response to *E. necator* at 1 dpi. The highest number of up-regulated genes associated with disease resistance at 1 dpi was detected in *Ren3*^+^, *Ren4D*^+^, and *Ren4U*^+^ vines (199, 193, and 167 genes, respectively), while a lower number of up-regulated genes was found in *Ren2*^+^, *Run1*^+^, *Run1.2b*^+^, *Run2.2*^+^ leaves (93, 59, 70, and 42 genes, respectively). The *Run2.1*^+^ accession was the PM-resistant with the fewest up-regulated genes associated with PM resistance, with only 7 and 14 genes at 1 and 5 dpi, respectively. Similar patterns were observed for the down-regulated genes detected only in PM-resistant vines.

### Overlap in transcriptional modulation associated with powdery mildew resistance between *R* loci

PM resistance-associated DEGs were compared across PM-resistant accessions to evaluate the overlap in defense responses between the different PM resistance loci (**Figure 5 & 6**; **Supplementary Figures 4-7**). On average, the *R* loci shared 52.9 ± 10.4 % and 38.0 ± 13.6 % of their defense-related DEGs associated with disease resistance with at least another *R* locus at 1 and 5 dpi, respectively. However, the overlap of PM resistance-associated DEGs between two *R* loci was limited (8.4 ± 10.0 %). The largest pairwise overlaps were found among the up-regulated genes at 1 dpi between the *Ren3*^+^, *Ren4D*^+^, and *Ren4U*^+^ genotypes (**Figure 5A, Supplementary Figure 4**). The *Ren3*^+^ and *Ren4D*^+^ accessions shared 64 up-regulated genes, while 49 and 44 defense-related genes were up-regulated in both *Ren3*^+^ and *Ren4U*^+^ genotypes, and both *Ren4D*^+^ and *Ren4U*^+^ vines, respectively. The overlap of up-regulated genes between *Ren3*^+^ and *Ren4D*^+^ accessions encompassed the largest number of extra- and intracellular receptor genes (15 RLK and 10 NLR genes), MAPK-signaling genes (7), and phytoalexin biosynthesis-related genes (11) including six stilbene synthase genes (**Figure 5A**). Eight genes involved in ROS production were found up-regulated in several *R* loci: a copper amine oxidase, three respiratory burst oxidase homolog (RBOH), and four class III cell wall peroxidase (Prx) genes. Regarding ROS scavenging, two catalase genes were up-regulated in both *Ren3*^+^, *Ren4D*^+^, and *Ren4U*^+^ leaves, as well as seven glutathione *S*-transferase genes in *Ren3*^+^ and *Ren4U*^+^ accessions. Some hormone-mediated signaling genes were also found up-regulated genes in several *R* loci at 1 dpi, including one gene involved in SA signaling, 12 genes in JA signaling, 13 genes in ET signaling, and 11 genes in ABA signaling. The shared JA-signaling genes encompassed 8 genes involved in JA biosynthesis, including an allene oxide synthetase (AOS) gene (VIT_18s0001g11630) that was more highly expressed in *Ren3*^+^, *Ren4D*^+^, *Ren4U*^+^, *Run1.2b*^+^ leaves in response to *E. necator*. The gene *VviJAZ4* (VIT_09s0002g00890) encoding a jasmonate-ZIM-domain protein was also up-regulated by the same PM resistance loci. Ectopic expression of *VqJAZ4* from *V. quinquangularis* was showed to enhance resistance to PM in *A. thaliana*, suggesting a role of the jasmonate-ZIM-domain protein in grape PM resistance (Zhang *et al*., 2019). Furthermore, *VviMYC2* (VIT_02s0012g01320) was up-regulated in *Ren2*^+^, *Ren3*^+^, *Ren4D*^+^, and *Ren4U*^+^ accessions at 1 dpi but not in any genotype possessing a *R* locus from *M. rotundifolia*. The same pattern was observed for *VviJAR1* (VIT_15s0046g01280) which was up-regulated only in *Ren3*^+^, *Ren4D*^+^, and *Ren4U*^+^ accessions. In *A. thaliana*, JAR1 encodes a jasmonate-amido synthetase that catalyzes the formation of the biologically active jasmonyl-isoleucine (JA-Ile) conjugate, while MYC2 is a basic helix-loop-helix (bHLH) transcription factor that is involved in JA signaling but also in ABA- and SA-mediated signaling (Abe *et al*., 2003; Dombrecht *et al*., 2007; Gautam *et al*., 2021; Staswick and Tiryaki, 2004). Regarding cell wall reinforcement, a dirigent protein-encoding gene (VIT_06s0004g01020) was up-regulated in *Ren4U*^+^, *Run1*^+^ and *Run2.2*^+^ genotypes at 1 dpi. Dirigent proteins participate in the biosynthesis of lignans and lignin which both play a role in plant defense. Lignans inhibit microbe-derived degradative enzymes while lignin accumulation in the plant cell wall forms a physical barrier against pathogens (Paniagua *et al*., 2017). In addition, the callose synthase genes *VviCalS1* and *VviCalS3* (Yu *et al*., 2016) were up-regulated in only *Ren4D*^+^ and *Ren4U*^+^ accessions at 1 dpi and 5 dpi, respectively (**Figures 5A & 6A, Supplementary Figures 4 & 6**). These results support previous observations which suggested that *Ren4*-mediated resistance relies on the encasement of the pathogen haustorium in callose in addition to PCD of the infected cells (Qiu *et al*., 2015).

**Figure 5:**
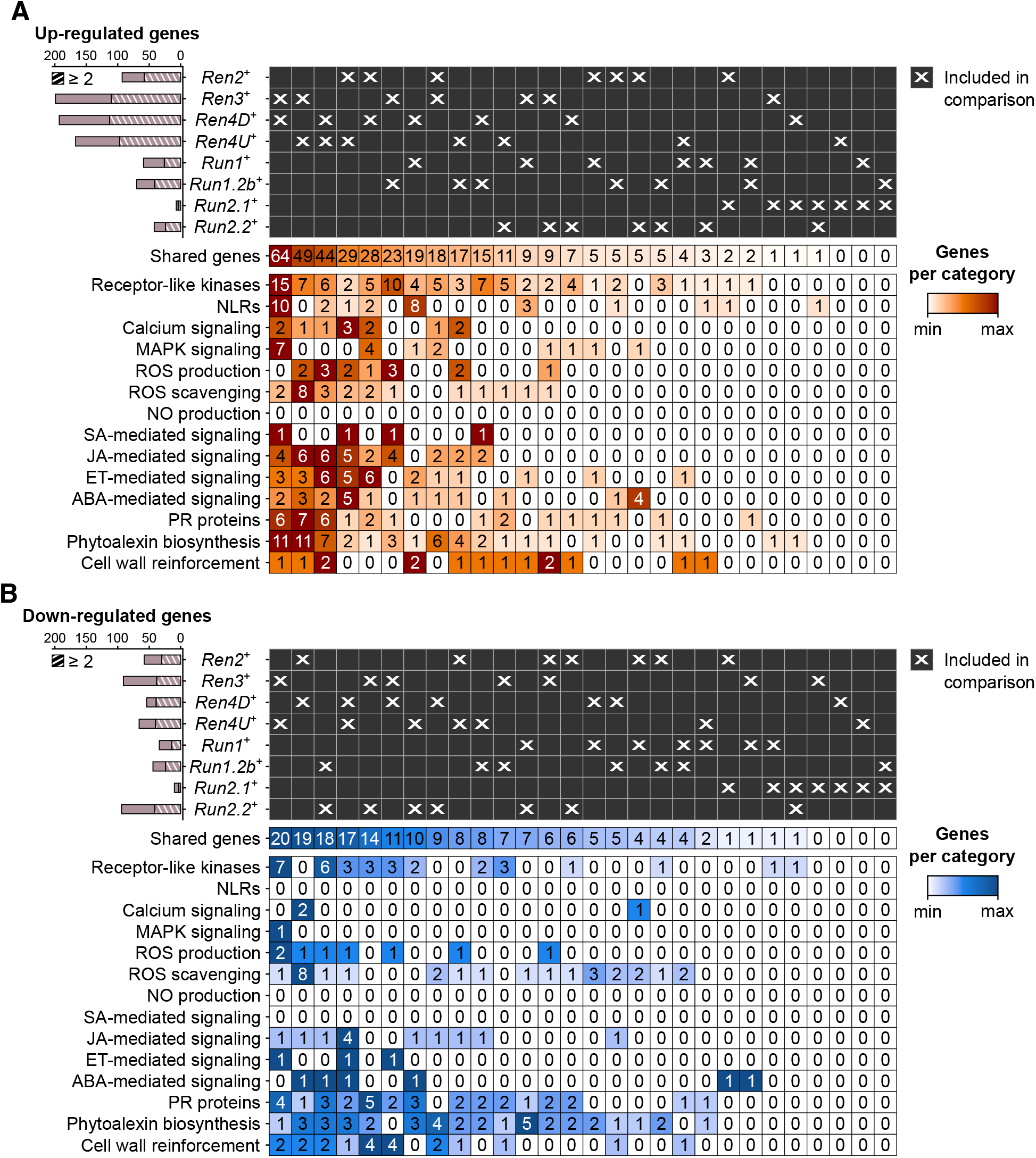
Pairwise comparison of the defense-related genes found up- and down-regulated in response to *E. necator* at 1 dpi in only the eight *R* loci. Striped bar plots depict the number of defense-related genes differentially expressed in at least two *R* loci.

Regarding the down-regulated genes associated with PM resistance at 1 dpi, *Ren3*^+^ and *Ren4U*^+^ accessions shared the highest number of genes with a lower gene expression in response to *E. necator* (20), followed by the *Ren2*^+^ and *Ren4D*^+^ vines (19), the *Run1.2b*^+^ and *Run2.2*^+^ plants (18), and the *Ren4D*^+^ and *Ren4U*^+^ genotypes (17; **Figure 5B**). The overlap in down-regulated genes between the *Ren3*^+^ and *Ren4U*^+^ accessions comprised 7 RLK genes, 2 Prx genes, and 2 cellulose synthase-like genes. Both *Ren2*^+^ and *Ren4D*^+^ leaves exhibited a lower gene expression of 8 ROS scavenging-related genes in response to *E. necator* at 1 dpi. Four fatty acid desaturase genes involved in JA biosynthesis were down-regulated in the *Ren4D*^+^ and *Ren4U*^+^ genotypes, and an ERF gene was down-regulated in *Ren3*^+^, *Ren4D*^+^ and *Ren4U*^+^ plants.

At 5 dpi, the highest overlap of DEGs associated with PM resistance was found between *Ren4U*^+^ and *Run1*^+^ genotypes, with 31 and 25 common up- and down-regulated genes, respectively (**Figure 6**). The shared up-regulated genes between *Ren4U*^+^ and *Run1*^+^ accessions encompassed 5 RLKs and 17 NLRs, 3 MAPKK genes, 3 genes involved in ABA signal transduction, and the gene encoding the exoribonuclease 4 (XRN4/EIN5; VIT_14s0030g01580) involved in ethylene response (**Figure 6A**). Five additional ethylene-signaling genes were found up-regulated in *Ren2*^+^ and *Ren4D*^+^ accessions at 5 dpi: a 1-aminocyclopropane-1-carboxylate synthase (ACS) gene (VIT_02s0025g00360) and four ERF genes. In contrast, the 25 down-regulated genes shared by *Ren4U*^+^ and *Run1*^+^ genotypes included six genes associated with ROS production (1) and scavenging (6), and 10 genes involved in the biosynthesis of anthocyanins and condensed tannins (**Figure 6B**).

**Figure 6:**
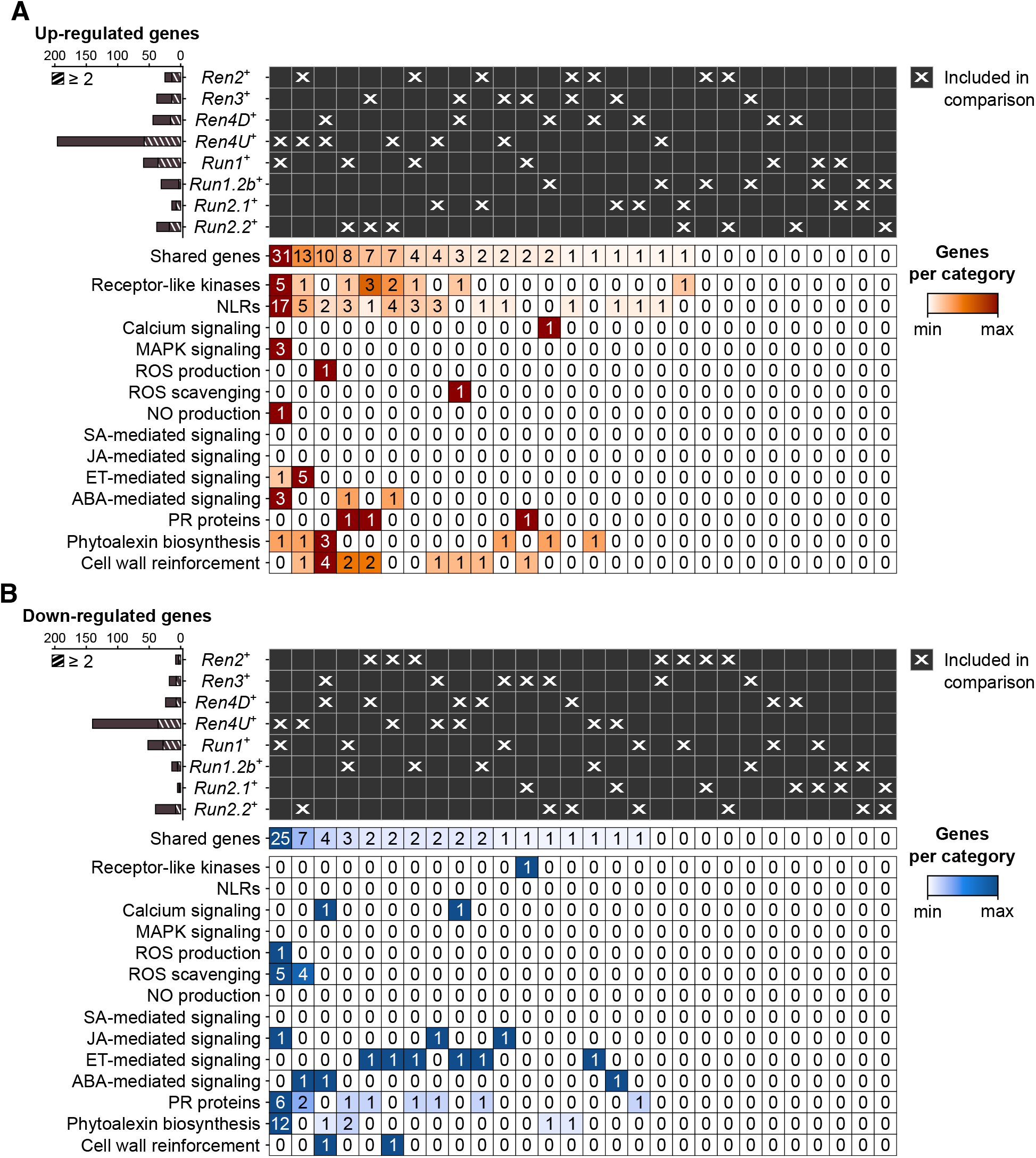
Pairwise comparison of the defense-associated genes found up- and down-regulated in response to *E. necator* at 5 dpi in only the eight accessions carrying a *R* locus. Striped bar plots represent the defense-related genes differentially expressed in at least two *R* loci.

### Resistance locus-specific transcriptional modulations in response to *E. necator*

Nearly half of the PM resistance-associated DEGs were found in only one PM resistance locus (47.1 ± 10.4 % and 62.0 ± 13.6 % at 1 and 5 dpi, respectively). Although the *Ren3*^+^, *Ren4D*^+^, and *Ren4U*^+^ genotypes exhibited the greatest overlap of PM resistance-associated up-regulated genes at 1 dpi, the three PM resistance loci also had the highest number of genes (89, 80, and 70 genes, respectively) that were up-regulated in a *R* locus-specific manner (**Figure 7**). Regarding the down-regulated genes at 1 dpi, *Ren3*^+^ and *Run1.2b*^+^ plants had the most numerous *R* locus-specific ones (53 genes each). At 5 dpi, the largest number of *R* locus-specific DEGs was identified in *Ren4U*^+^ leaves, with 138 up-regulated and 104 down-regulated genes.

**Figure 7:**
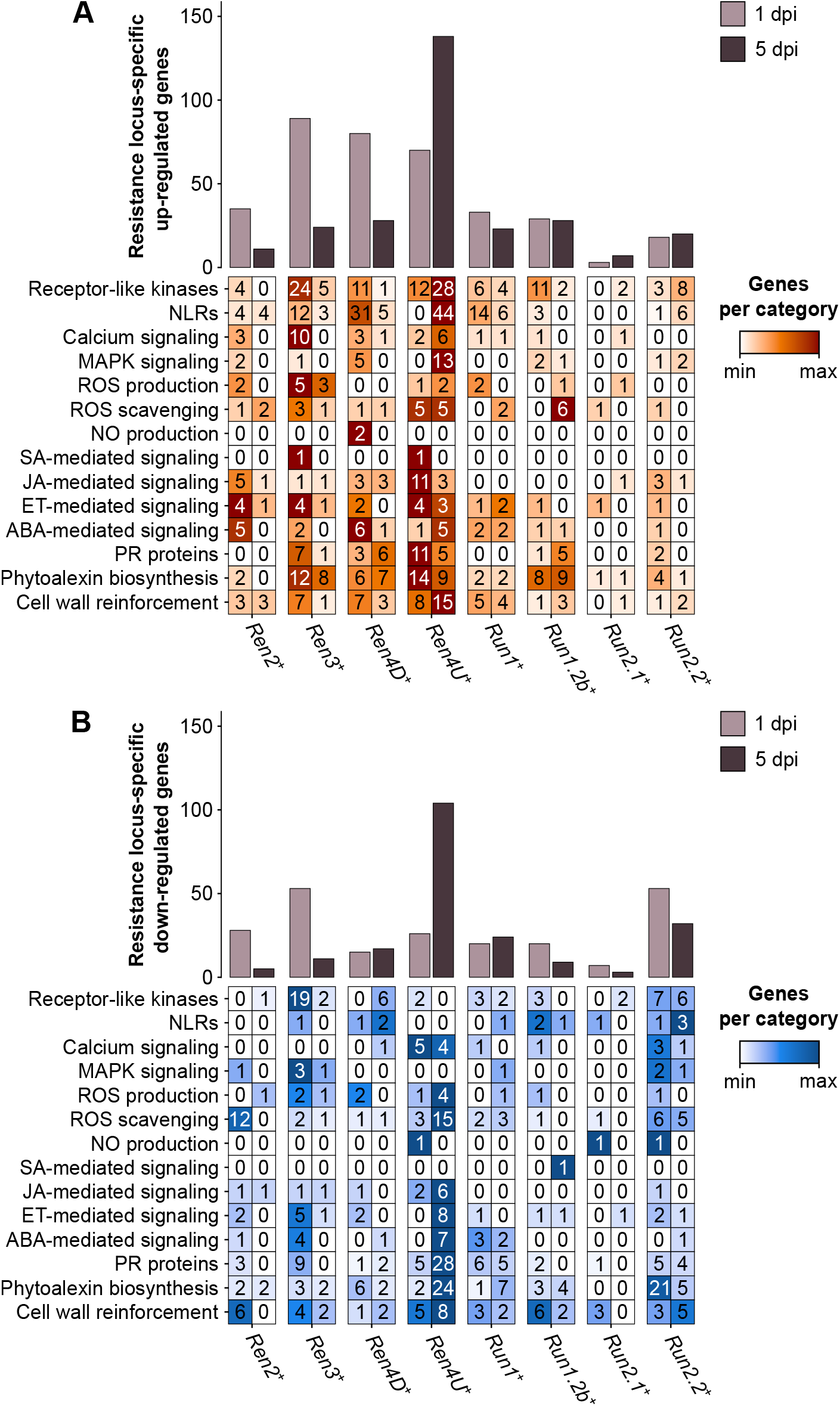
Defense-related categories among the genes identified as up-regulated (**A**) and down-regulated (**B**) by only one single PM resistance locus in response to *E. necator*.

Differences in defense-related categories among the *R* locus-specific DEGs could be observed between the eight *R* loci and the two time points (**Figure 7**). The *Ren3*^+^ accession exhibited the greatest number of *R* locus-specific DEGs encoding intracellular receptors (RLKs) at 1 dpi, with 24 up-regulated and 19 down-regulated genes. In contrast, the presence of *E. necator* led to the up-regulation of 28 RLK and 44 NLR genes in only the *Ren4U*^+^ leaves at 5 dpi. In addition to the intra- and extracellular receptors, the *R* locus-specific DEGs encompassed several PR protein genes as well as genes involved in phytoalexin biosynthesis and cell wall rearrangement. Regarding the PR protein genes, the *Ren4U*^+^ accession showed the largest number of up-regulated genes at 1 dpi (11) but also the most numerous down-regulated genes at 5 dpi (28). The *Ren3*^+^ genotype had also some *R* locus-specific DEGs encoding PR proteins at 1 dpi (7 up-regulated and 9 down-regulated genes), while the *Ren4D*^+^ and *Run1.2b*^+^ vines had six and five specific up-regulated ones at 5 dpi, respectively. The PM resistance locus-specific DEGs encoding PR proteins included PR1-like proteins, beta-1,3-glucanases (PR2), basic chitinases (PR3), thaumatin-like proteins (PR5), subtilisin-like endoproteinases (PR7), chitinases type I (PR11) and III (PR8), Bet v 1 homologs (PR10), lipid-transfer proteins (PR14), and germin-like proteins (PR16). PM resistance loci also exhibited differences in *R* locus-specific DEGs involved in the biosynthesis of antimicrobial compounds. Like the PR proteins, the *Ren4U*^+^ accession had the highest number of genes that were up-regulated at 1 dpi (14) and down-regulated at 5 dpi (24) in a PM resistance locus-specific way. In addition to the *Ren4U*^+^ accession, several phytoalexin biosynthesis-related genes were found up-regulated only in the *Ren3*^+^, *Ren4D*^+^, and *Run1.2b*^+^ genotypes at both time points. In contrast, 21 genes involved in the biosynthesis of phytoalexins were found down-regulated specifically in the *Run2.2*^+^ plants at 1 dpi, including 18 genes from the phenylpropanoid biosynthesis pathway. The *Ren3*^+^-specific up-regulated genes at 1 dpi comprised three alkaloid-related genes (berberine bridge enzymes), three genes from the phenylpropanoid biosynthesis pathway: two cinnamate 4-hydroxylases (C4H) and one 4-coumarate--CoA ligase (4CL), and two stilbene synthases. Genes involved in the biosynthesis of terpenoids and triterpenoids were also found up-regulated in only one *R* locus at 1 dpi, such as two and three terpene synthase genes in *Ren4D*^+^ and *Ren4U*^+^ accessions, respectively, as well as two and three oxidosqualene cyclase genes in *Run2.2*^+^ and *Ren4D*^+^ vines, respectively. Concerning the *R* locus-specific DEGs involved in cell wall rearrangement, the *Ren4U*^+^ genotype had the largest number of both up- and down-regulated genes. The *Ren4U*^+^-specific up-regulated genes at 5 dpi included five cellulose synthase genes, six cellulose synthase-like genes and two callose synthase genes: *VviCalS7* (VIT_12s0028g00400) and *VviCalS10* (VIT_17s0000g10010). Another callose synthase gene, *VviCalS11* (VIT_00s0265g00050), was also found up-regulated in *Run1*^+^ plants at 5 dpi. This result suggests that *VviCalS11* could play a role in the accumulation of callose deposits described in *Run1*^+^ plants (Feechan *et al*., 2015).

## Discussion

### Powdery mildew resistance loci confer different levels of resistance to *E. necator*

Evaluation of PM development using microscopy, visual scoring, and total *E. necator* transcript abundance, revealed differences of intensity and timing of the response to *E. necator* at 5 and 14 dpi (**Figure 1**; **Supplementary Figure 1**). In particular, the *Ren2, Ren3* and *Run2.2* loci conferred a lower level of resistance to PM compared to the five other *R* loci. Similar disease phenotypes have been described for these *R* loci in previous studies (Qiu *et al*., 2015; Dry *et al*., 2019). The absence of primary hypha on the leaves of the *Ren4D*^*+*^, *Ren4U*^*+*^, *Run1*^*+*^, and *Run1.2b*^+^ accessions and the very low transcript abundance for *E. necator* at 5 dpi suggest that restraint of the pathogen growth occurs rapidly in these vines, likely after haustorium formation at approximately 1 dpi (Leinhos *et al*., 1997). In contrast, the presence of some secondary and tertiary hyphae on *Ren2*^+^, *Ren3*^+^, *Run2.1*^+^, and *Run2.2*^+^ genotypes’ leaves suggest that restriction of *E. necator* happens at later time, between 1 and 2 dpi (Leinhos *et al*., 1997). In addition, fungal structures covered most of the leaves of *Ren2*^+^ and *Ren3*^+^ vines at 14 dpi, suggesting that these two *R* loci are less efficient in restricting the pathogen growth. Assessing the development of *E. necator* at additional time points post-inoculation and monitoring PCD would help to narrow down the difference of timing in the response to PM between the *R* loci. Furthermore, repeating the disease evaluation with additional breeding lines for each *R* locus would allow to confirm the observed phenotypes; while repeating the inoculation with additional *E. necator* isolates would help determine if the observed phenotypes are strain-specific.

### Adapting the reference transcriptome allows comparing the transcriptional modulations of defense-related genes between different *Vitis* backgrounds

This study encompassed fourteen accessions with different genetic backgrounds making the comparison of the gene expression from orthologous genes challenging. To cope with this challenge, we incorporated small sequence polymorphisms (SNPs and INDELs) compared to the grape PN40024 CDS into a diploid synthetic transcriptome for each grape accession and performed a *de novo* assembly of the unaligned RNA-seq reads. The sequence variant analysis revealed an elevated number of heterozygous SNPs in the leaf transcriptomes of interspecific and intergeneric hybrids compared to pure *V. vinifera* cultivars, reflecting the genetic diversity among *Vitis* species and between *M. rotundifolia* and *Vitis* genera. The construction of the fourteen accession-specific reference transcriptomes improved the alignment of the RNA-seq reads (**Figure 3C**). Complete genome and/or transcriptome references of each grape accession would provide a more comprehensive representation of the defense-related genes and potentially a more accurate sequence for read alignment.

To assess the functional overlap among defense mechanisms between PM resistance loci, we focused our study on the transcriptional modulation of 2,694 defense-related genes in response to PM. These genes were selected based on functional domain composition and sequence similarity with proteins of *A. thaliana*. Although the defense-related genes identified might not be exhaustive and not definitive, the refinement of the functional annotation of PN40024 that was performed in this study represents a valuable resource for the present and future transcriptomic studies of grape-microbe interactions.

### Powdery mildew resistance loci trigger the transcriptional modulation of different defense-related genes in response to *E. necator*

By comparing the transcriptomes of PM- and mock-inoculated leaves of eight PM-resistant and six PM-susceptible grape accessions, we identified the defense-related DEGs associated with disease resistance (**Figure 4**). Although the comparison of the PM resistance-associated DEGs showed that the *R* loci shared DEGs to some extent, the overlap between two *R* loci was restricted (8.4 ± 10.0 %). It is worth noting that no defense-related gene was found differentially expressed in response to *E. necator* in more than five PM resistance loci (**Supplementary Figures 4-7**). In addition, around half of the PM resistance-associated DEGs of each *R* locus was found specific (**Figures 5 & 6**). Comparison of the *R* locus-specific DEGs showed that the eight *R* loci extensively differ in their transcriptional response to *E. necator* through the transcriptional modulation of specific defense-related genes (**Figure 7**). Interestingly, different allelic forms of the *Ren4, Run1*, and *Run2* genetic loci exhibited different responses to *E. necator* and even shared a larger number of DEGs with another genetic locus rather than with their respective haplotype (**Figures 5 & 6**).

To our knowledge, this study corresponds to the first attempt of comparing the transcriptional modulations in response to PM between multiple *R* genetic loci. Previous studies focused on a single *R* locus, a single PM-resistant accession (Fung *et al*., 2007; Jiao *et al*., 2021; Weng *et al*., 2014), or multiple haplotypes of a single genetic locus (Amrine *et al*., 2015). Comparisons with previous studies are difficult because different protocol of inoculation and different strains of *E. necator* were used. Repeating the transcriptome profiling of additional recombinant lines for each *R* locus, both PM-resistant and PM-susceptible, combined with whole-genome sequencing would help dissect the *R* loci and identify the genes essential for resistance. Including other time points may provide additional information to understand gene expression modulation earlier and later than what considered in this study. For instance, few DEGs were detected for the *Run2.1*^+^ accession, suggesting either that PM resistance is associated with the expression modulation of few genes or that the transcriptional reprogramming in response to *E. necator* occurs at a different time for the *Run2.1* locus.

Finally, this dataset represents a valuable resource for future breeding perspectives as it could help select the most functionally diverse *R* loci to introgress into *V. vinifera*.

## Data availability statement

RNA sequencing data are accessible through NCBI under the BioProject PRJNA897013.

## Author contributions

D.C, M.A.W, S.R, and M.M. designed the project. S.R and D.P performed the sample inoculation, collection, microscopy, and visual scoring. R.F.-B. extracted RNA and prepared sequencing libraries. M.M. performed the data analyses. M.M and D.C wrote the manuscript.

